# Unconventional cell division cycles from marine-derived yeasts

**DOI:** 10.1101/657254

**Authors:** Lorna M.Y. Mitchison-Field, José M. Vargas-Muñiz, Benjamin M. Stormo, Ellysa J.D. Vogt, Sarah Van Dierdonck, Christoph Ehrlich, Daniel J. Lew, Christine M. Field, Amy S. Gladfelter

## Abstract

Fungi have been found in every marine habitat that has been explored, however, the diversity and functions of fungi in the ocean are poorly understood. In this study, fungi were cultured from the marine environment in the vicinity of Woods Hole, MA, USA including from plankton, sponge and coral. Our sampling resulted in 36 unique species across 20 genera. We observed many isolates by time-lapse differential interference contrast (DIC) microscopy and analyzed modes of growth and division. Several black yeasts displayed highly unconventional cell division cycles compared to those of traditional model yeast systems. Black yeasts have been found in habitats inhospitable to other life and are known for halotolerance, virulence, and stress-resistance. We find that this group of yeasts also shows remarkable plasticity in terms of cell size control, modes of cell division, and cell polarity. Unexpected behaviors include division through a combination of fission and budding, production of multiple simultaneous buds, and cell division by sequential orthogonal septations. These marine-derived yeasts reveal alternative mechanisms for cell division cycles that seem likely to expand the repertoire of rules established from classic model system yeasts.

## Introduction

Fungi are critical components of the biosphere with diverse roles in cycling nutrients, shaping microorganism communities, and acting as opportunistic pathogens. These key functions have been extensively explored in terrestrial ecosystems, but the roles of fungi in the marine environment are much less appreciated. Large-scale expeditions sampling marine microbiological diversity would have largely missed fungi for technical reasons such as the use of size filtration and the limitations of markers for systematic molecular identification of fungi [1]. Nevertheless, fungi have been found in every part of the marine environment where they have been investigated, associated with marine sediments, invertebrates, marine mammals, algae, driftwood, and throughout the water column [2–13]. Thus, there is major untapped and unknown biodiversity of this major kingdom of life in the oceans [14].

Phylogenetics studies point to a terrestrial origin of multiple obligate marine fungal lineages [15]. This transition from terrestrial to marine environments may have occurred multiple times, as many of the fungi detected in the marine environments have been previously described in terrestrial habitats [16–18]. However, these marine-derived fungi were detected in samples collected far from shore, suggesting that they are actual inhabitants of the aquatic environment rather than recent arrivals from land. Some of these fungi seem to have a truly amphibious lifestyle, based on gene expression data [19] and strong correlations with abiotic factors [20, 21]. This is indicative of the adaptability of the fungal kingdom and makes it challenging to define what truly constitutes a marine (as opposed to terrestrial) fungus. Suggested criteria include that the fungus was isolated from marine environments on multiple occasions, can grow on marine-origin substrates, forms ecologically-relevant relationships with other marine organisms (pathogen, symbiont, etc.), and has adapted to the marine environment as evident from genetic analyses or metabolic activity [2, 12, 13, 22, 23].

Our goal was to identify culturable species of fungi from the sea to assess how fungi in non-terrestrial environments grow and divide. Fungi in these settings face a myriad of potential stresses from temperature, high salinity, buoyancy challenges, UV exposure, and limited organic matter for nutrients. A subgroup of fungi of particular interest are the melanized fungi, also known as black yeasts, that are in the class Dothideomycetes [24]. Black yeasts have attracted the attention of researchers due to their biotechnological potential, high stress tolerance, and ability to cause severe mycosis. Black yeasts have not only been identified in marine environments but also extreme habitats such as salterns, rocks, ice, and desert mats [25]. We speculated that their well-appreciated adaptability might reflect an expanded repertoire of mechanisms regulating growth and division cycles.

Much of our current understanding of the cell division cycle derives from classic studies originating in the 1970s in two model yeast systems: the budding yeast *Saccharomyces cerevisiae* and the fission yeast *Schizosaccharomyces pombe* [26–31]. These evolutionarily-distant yeast species were intensively mined for cell division cycle mutations and, together with work from marine invertebrate embryos, gave rise to the molecular framework for understanding the cell cycle that we have today. In this framework, a cell-cycle network with properties of a relaxation oscillator (positive feedback, bistability, and delayed negative feedback) controls the synthesis and degradation of cyclins and other regulators of a family of cyclin-dependent kinases (CDKs) and phosphatases. These in turn trigger cell-cycle events through phosphorylation/dephosphorylation of target substrates. In addition, a series of checkpoint pathways can stop the oscillator as needed when extra time is required to avoid errors or in response to external signals or nutrients. Cell-cycle progression is regulated so as to maintain a consistent distribution of cell sizes in the population, although the mechanisms underlying such size control remain contested. In addition to size control, budding yeast cells have the added challenge of coordinating bud formation and mitosis so that daughter cells receive a nucleus, and these bud-bound nuclei must navigate the narrow passage of the mother-bud neck. Tight controls have been identified that work to ensure that only a single bud is produced and to sense if a bud has been built [32–34].

Despite sharing many common core components, notable contrasts have emerged between the two model yeast systems. There are substantial differences in how these morphologically distinct yeasts regulate the cell cycle, including how cell size is measured, how nutrient conditions regulate mating and meiosis [35–37], how the substrate specificity of the CDK is controlled [38–40], and how polarity and cytokinesis are regulated by the cell cycle [34]. These variations may not fully capture the variability in mechanisms controlling division. In this study, we analyze four different black yeasts isolated from marine environments and find that the division cycles of these yeasts display far more plasticity and behave in unconventional ways not predicted by the studies from model yeasts.

## Results

### Fungal isolation from the marine environment

Between 2016-2018, we sampled locations around Woods Hole, MA to culture fungi from the marine environment. Fungi were isolated from plankton tows of ocean water in Buzzards Bay and Martha’s Vineyard Sound, sediment from local beaches, a salt marsh, the sponge, *Cliona celata* and the coral, *Astrangia poculata* (Figure 1, Table 1). The two bodies of water are physically separated by the southwest corner of mainland Cape Cod and the Elizabeth Islands. Buzzards Bay is fed by the Acushnet River via the New Bedford Harbor, an 18,000-acre Environmental Protection Agency (EPA) superfund site. New Bedford Harbor has been the focus of restoration projects since the EPA found high concentrations of polychlorinated biphenyls (PCBs) and heavy metals such as cadmium and lead in the mid 1970’s [41]. Martha’s Vineyard Sound is encompassed by mainland Cape Cod, Martha’s Vineyard, and Nantucket, but still experiences strong tides and currents.

**Figure 1.**
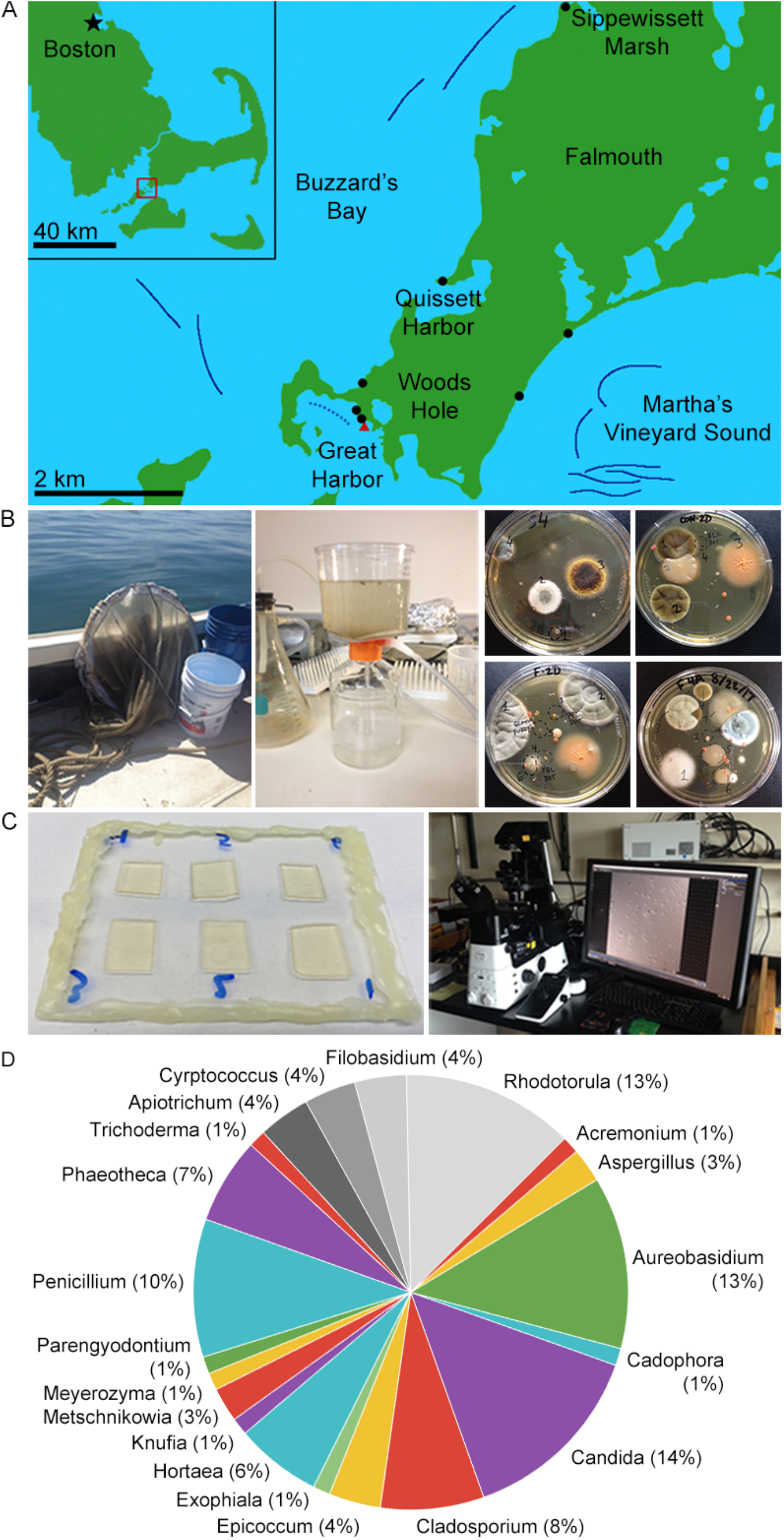
Culturing marine fungi from marine and coastal environments around Woods Hole, MA. A. Collection sites. Solid dark blue lines denote cruise tracks of plankton tows from summers of 2017 and 2018. The dotted blue line denotes the approximate cruise track of a Great Harbor tow conducted days after a small oil spill. Black dots mark beach sediment samples and the red triangle marks where coral and sponge were collected, however the only sediment samples sequenced so far have been from Sippewissett Marsh. B. Collection and culture methodology. Water column samples were collected by conducting plankton tows 1-1.2 km offshore using a 504 μm mesh net (left). 1.0 L water was concentrated using a 22 μm filter (center). Cultures were grown on various media for 1-2 weeks at 18-20 °C. C. DIC imaging methodology. Samples were prepared six on a slide on agar pads of the same type on which they were cultured. Images were collected at 5 min intervals. We observed growth morphology by DIC time-lapse microscopy using a Nikon Eclipse Ti2 inverted microscope. D. Genus abundance. Numeric labels indicate number of isolates found belonging each of the 20 different genera we found. We found 16 different species of Ascomycota (in color) and 4 Basidiomycota (grayscale).

**Table 1.**
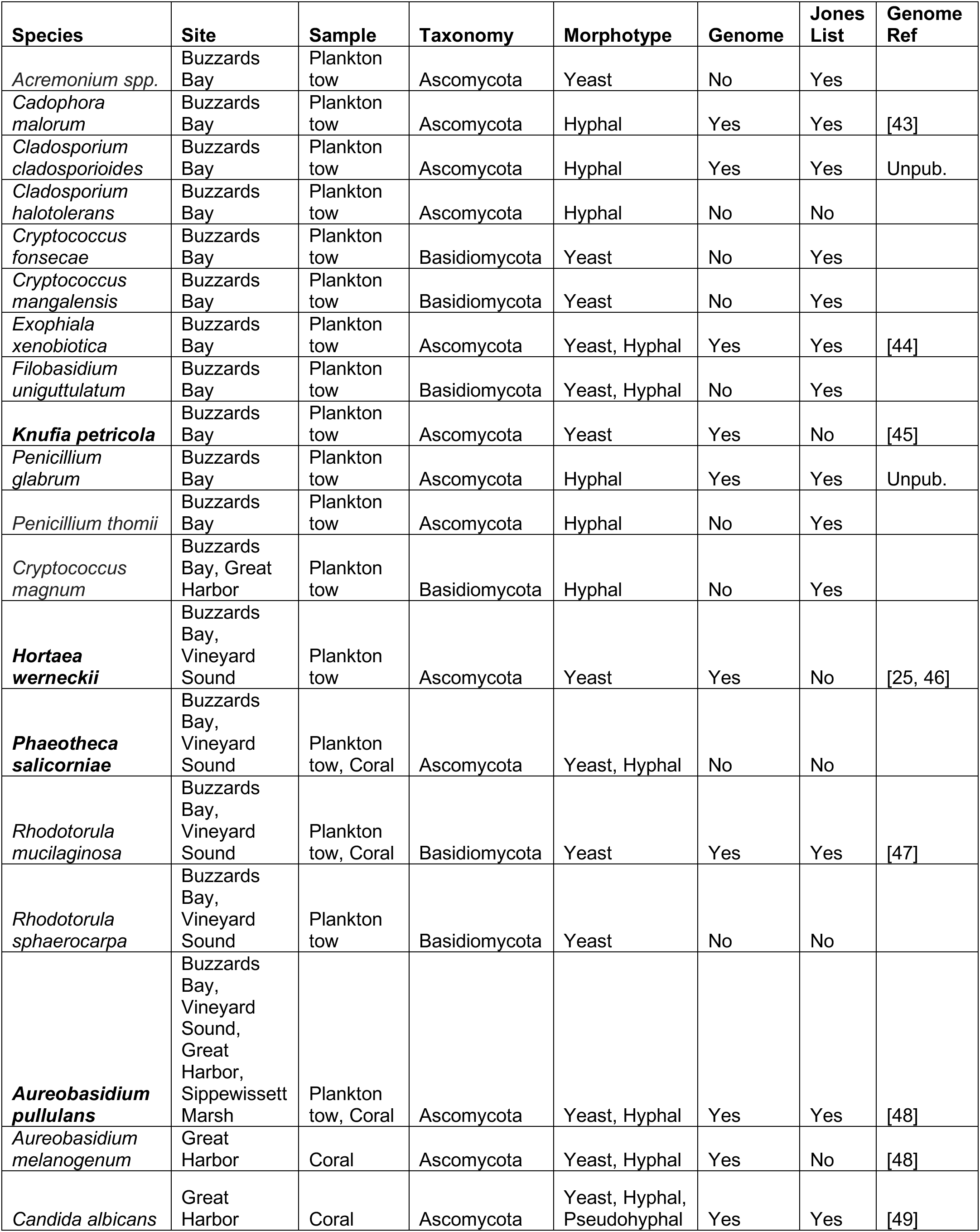
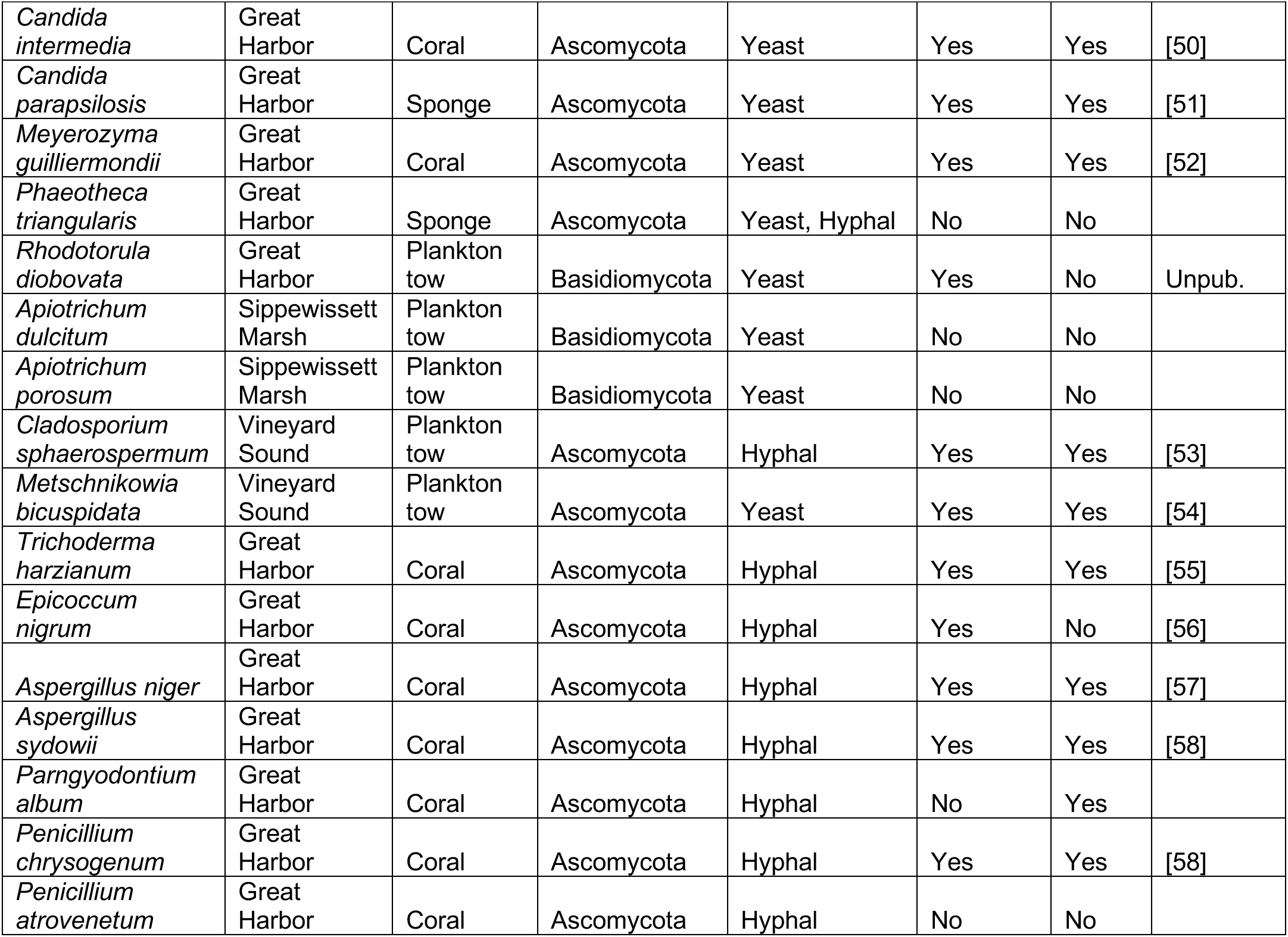
Fungal species isolated from marine environments near Woods Hole, Massachusetts.

After collection, samples were plated onto standard fungal media (with dextrose, malt or potato as carbon sources) made with sterile seawater and containing antibiotics to limit bacterial growth. Unconcentrated sea water collected in the vicinity of the samples was used as a control to ensure that isolates were not contaminants in the sampling pipeline. Individual colonies arising from plankton tows, sediment, sponge, or coral were then subcultured on individual plates, observed with DIC microscopy, and identified based on amplifying and sequencing the rDNA locus (Figure 1B-C, Table S1). We identified 36 species, including 16 Ascomycetes and 4 Basidiomycetes; 23 of the species were previously isolated from marine environments [42] (Figure 1D, Table 1). Multiple species were found uniquely in only one of the habitats sampled; others appeared across several collection sites.

The isolates exhibit a diversity of colony morphologies and pigmentation and include yeasts and filamentous fungi (Figure 2A). To assess morphogenesis and division patterns at the level of single-cells, a subset of the species were filmed using high-magnification, DIC time-lapse microscopy. This single-cell analysis revealed many modes of growth and division in the Woods Hole marine fungi collection including budding, fission, filamentous growth and meristematic growth. Here, we focus on four black yeasts that are morphologically distinct from each other and from model yeasts (Figure 2B-C). Three of these species display clear black colony growth with a tendency to produce terracotta pigments at the periphery of colonies, and one is initially pale pink and slowly melanizes over time under more stressed growth conditions. They each display surprising and unconventional division cycles.

**Figure 2.**
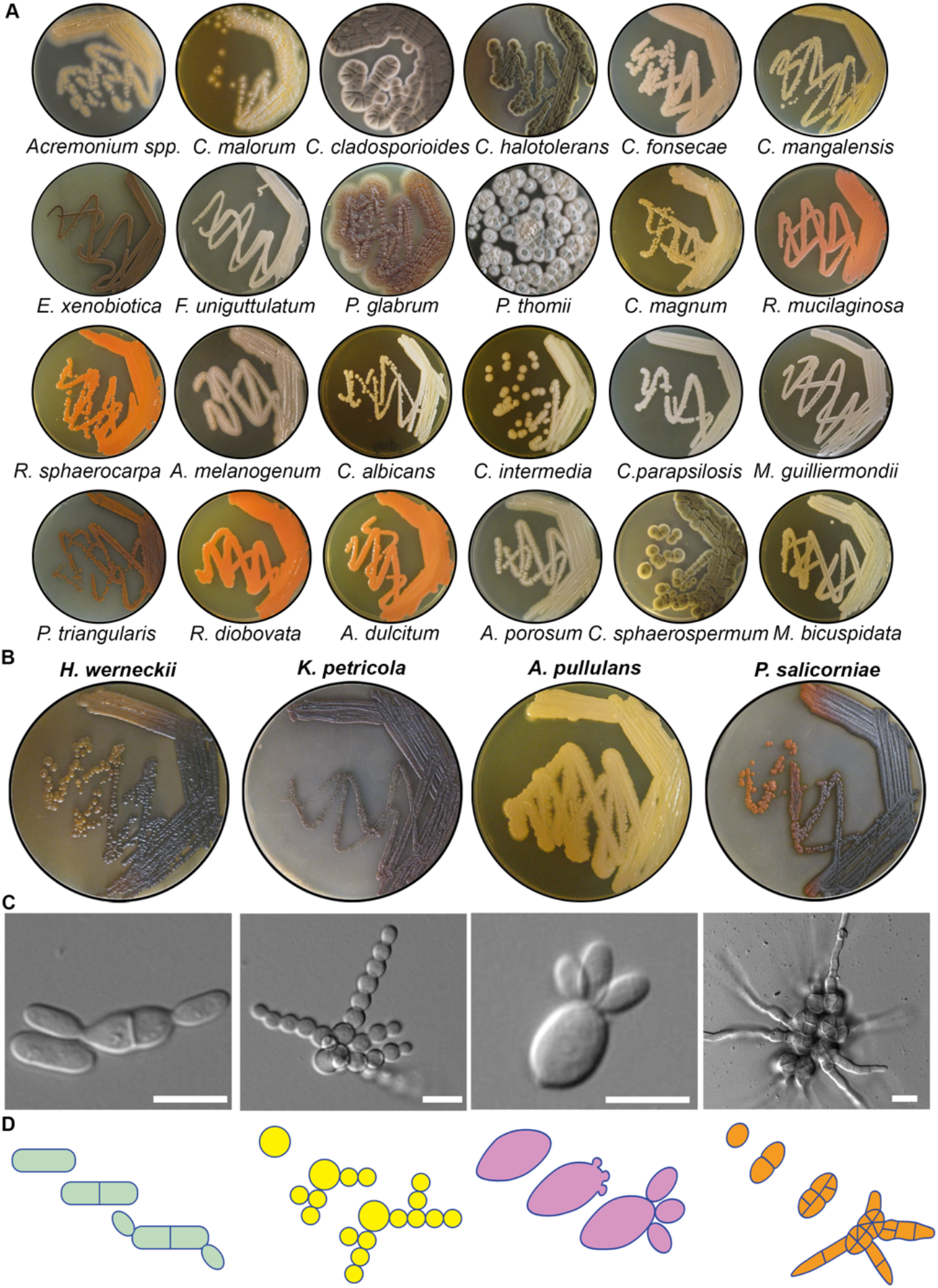
Overview of cultured marine-derived fungal diversity. A. Macroscopic colony morphology of identified marine fungi grown for 10 days on YPD + 0.6M NaCl. Images are ordered the same as in Table 1. B. Macroscopic colony morphology of the four black yeasts chosen for further characterization. C. Representative DIC images of each species showing division patterns. Scale bar, 10 μm. D. Schematic diagrams of colony growth for each species.

### Alternation between fission and budding in *Hortaea werneckii*

*H. werneckii* was identified in multiple locations and has been previously studied due to its extreme halotolerance with growth possible in up to 5 M NaCl [59]. Additionally, H. werneckii is the etiologic agent of tinea nigra, which is an asymptomatic superficial mycosis that is restricted to the palms of the hands and soles of the feet [60]. Remarkably, *H. werneckii* cells could divide both by fission and budding, with each cell compartment inheriting a single nucleus (Figure 3A-B, Video S1). After placement on a solid medium, cells grew both isotropically and from the cell poles and divided by development of medial septa. This is reminiscent of a typical fission yeast cell cycle with the exception that we did not observe separation of the two daughter cells after the septum was built. Remarkably, 92% of the next division cycles involved the production of a bud, with only 8% of cells displaying a second fission-like process (Figure 3C). Even those cells that performed a second fission then proceeded to bud in the third cell cycle, indicating that all cells switch division modes. There was a strong bias toward alternating between fission and budding rather than consistently dividing by one mode.

**Figure 3.**
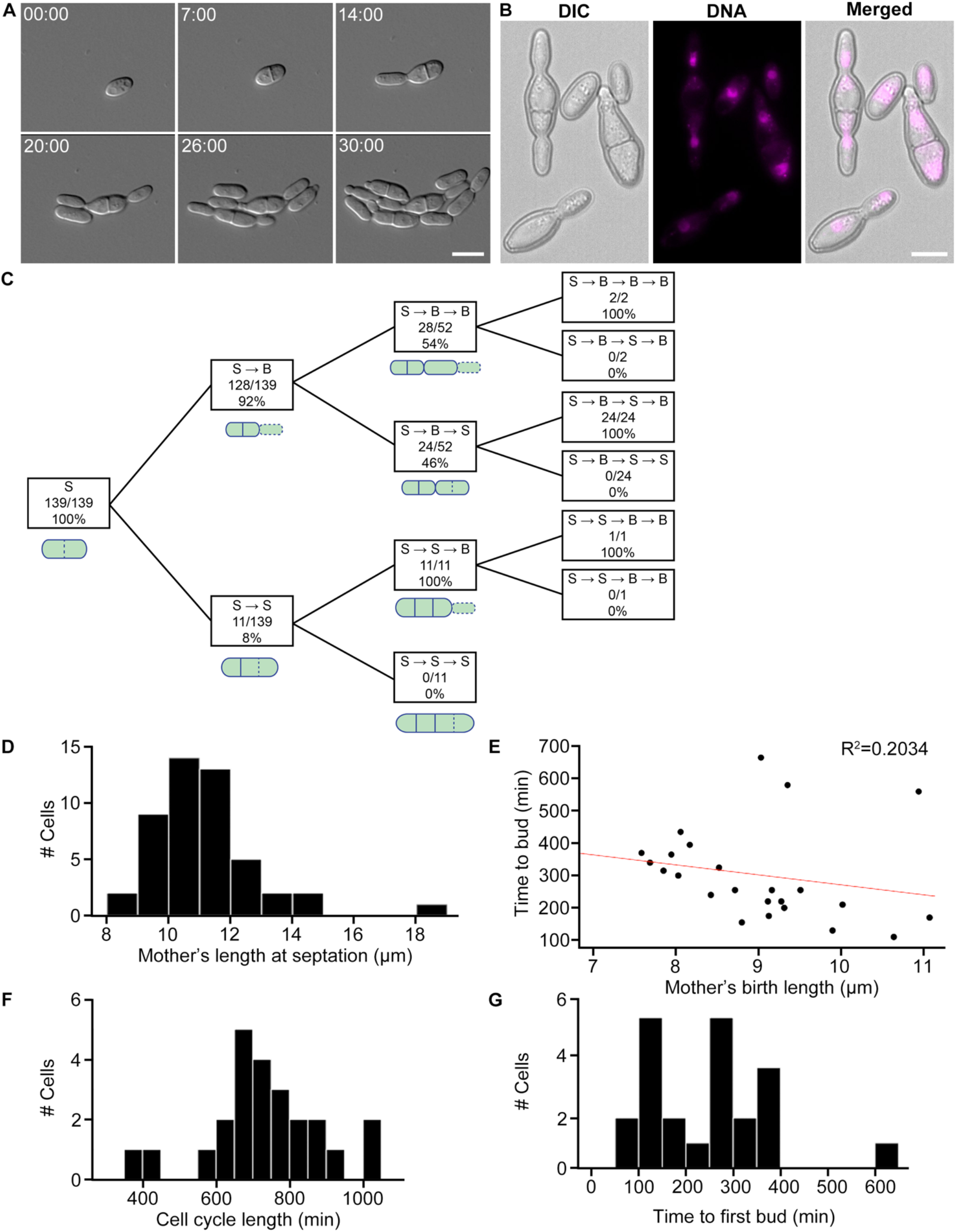
*Hortaea werneckii* cells divide via septation and budding. A. *H. werneckii* cell growth and development under 40x DIC. Scale bar, 10 μm. Time, Hours:Minutes. B. DNA staining in fixed cells. *H. werneckii* cells contain one nucleus per compartment. Mitosis seems to occur across the axis of cell division. Nuclei were stained using Hoechst. Scale bar, 10 μm. C. Quantifying sequence of cell division events. In this strain of *H. werneckii* we observed two cell division event types, budding and septation. Septation and budding events are represented by “S” and “B”, respectively. Dotted lines in the accompanying schematics indicate the latest division event type in that sequence. The diagram is interpreted as follows: of 139 total cells observed, all underwent septation as a first division event. Of those cells, 128 budded next (S → B) and 11 septated again (S → B). 52 of the 128 S → B cells divided again, 28 of which budded (S → B → B) and 24 septated (S → B → S). D. Mother’s length at septation. Mothers grew to an average length of 11.2 μm before septating (N = 48, SD = 1.7, CV = 15.2%). E. Mother’s birth length (μm) vs. cell cycle length (min). Birth length is defined as when the cell breaks off from its mother. Red line denotes linear fit. Larger cells have a shorter division time (N = 24, R^2^ = 0.203). F. Cell cycle length of *H. werneckii* (min). This is a measurement of the full cell cycle duration (N = 24, mean = 730, SD = 158, CV = 21.6%). G. Time to first bud (min). Time until a mother cell begins producing its first daughter cell (N = 24, mean = 253, SD = 124, CV = 49.0%).

Cell size and division times were quite variable. The median length at fission was 10.9 μm (N = 48, mean = 11.2 μm, standard deviation [SD] = 1.7 μm, coefficient of variation [CV] = 15.2%, Figure 3D), compared to Fission yeasts are known for their highly robust size control mechanisms, with mean length of 14 μm and CV of 7.7 in *S. pombe* [37]. Thus, *H. werneckii* shows substantially greater heterogeneity. The division site was generally centered in the cell (mean ratio of septated halves’ lengths = 1.02, SD = 0.15, CV = 14.2%, N = 48 pairs of septated halves). The median cell cycle length was 710 min (N = 24, mean = 730 min, SD = 158 min, CV = 21.6%, Figure 3E) for the cells undergoing medial septation. Plotting this time against size at birth did not reveal a strong negative correlation (Figure 3E), indicating no evidence of size control in *H. werneckii*. For budding cells, the median time for a bud to produce a new bud was 265 min (N = 24, mean = 253 min, SD = 124 min, CV = 49.0, Figure 3F), which is far more variable than for the model budding yeast *S. cerevisiae*. Thus, *H. werneckii* divides by fission and budding, generally alternating between modes of division from cell cycle to cell cycle. The fact that these distinct division patterns co-occur within the same organism indicates that they need not derive from extensive divergent evolution. In both modes, the cell cycle durations and cell sizes were far more variable in *H. werneckii* than in the model yeasts, and there was no obvious size control, indicating a high degree of plasticity in the growth and division programs.

### *Spherical* non-random budding produces dendritic morphologies in *Knufia petricola*

We isolated *K. petricola* from a plankton tow in Buzzards Bay. *K. petricola* is a melanized microcolonial fungus (MCF). MCF exhibit compact colonial structures and protective, melanized cell walls. They are extremotolerant and can grow on bare-rock surfaces. In addition to the Woods Hole marine environment, *K. petricola* has been found on rocks in extreme conditions from the Antarctic to dry deserts [61, 62] and as part of lichens [63]. This organism produces nearly spherical buds (Figure 4A). The next budding event is frequently opposite the previous division site (Figure 4B). Given how spherical these cells are, it is unclear how subsequent buds are so precisely positioned at the opposite side from the division site. After multiple cycles, chains of spherical cells are produced and at some frequency a chain will branch and produce a second chain with the colony taking on a dendritic-like shape (Figure 4A, Video S2).

**Figure 4.**
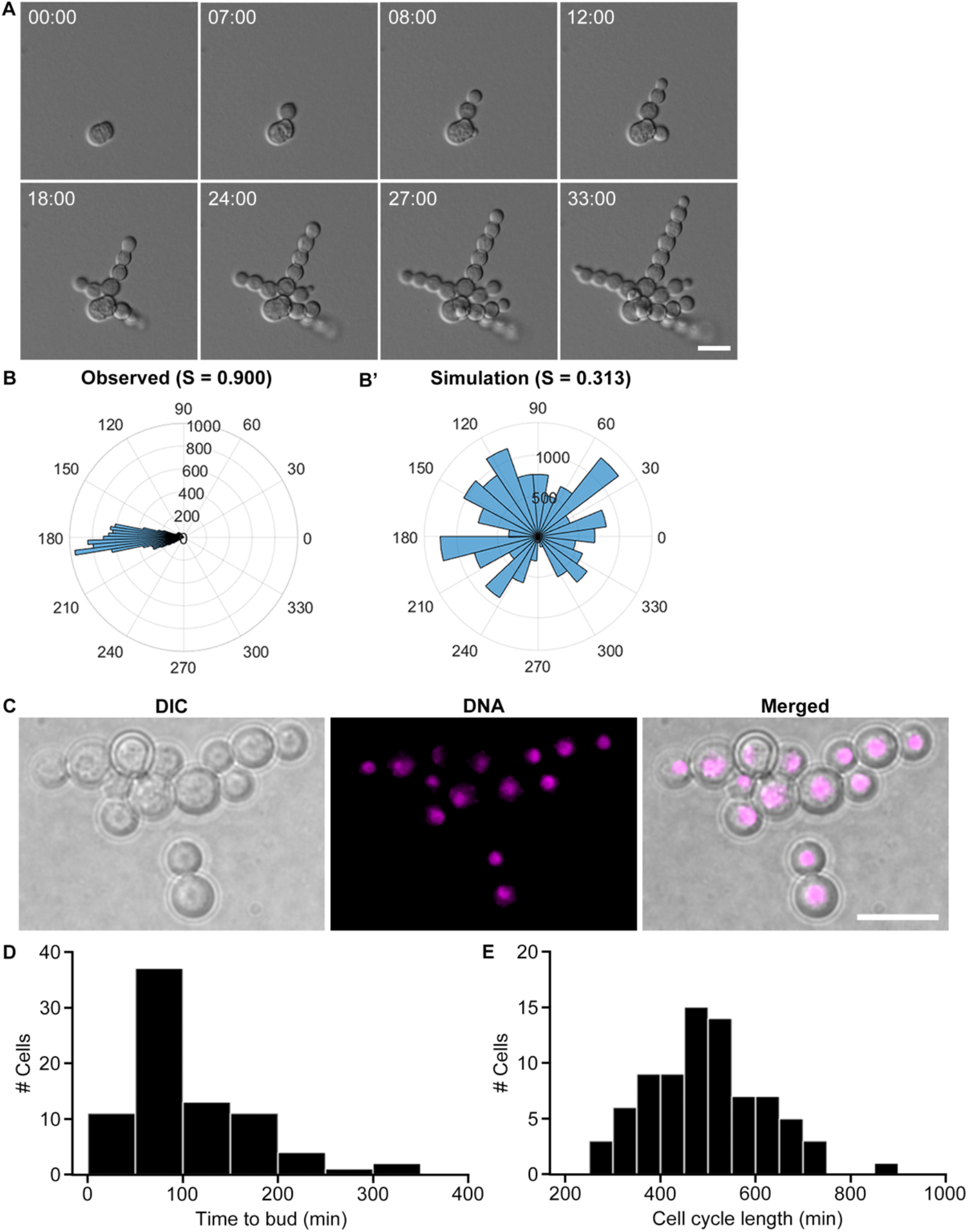
*K. petricola* growth through highly polarized patterning of spherical buds. A. Representative DIC images of *K. petricola* colony growth and development (40x). Scale bar, 10 μm. Time, Hours:Minutes. B. Observed angular distribution of colony growth. For every frame in the movie we quantified the average direction over which mass was added to each growing bud, where 0 degrees is defined at the site of the of old mother-bud connection. Note that growth is highly directional away from the old bud site, so it peaks at 180 degrees. Daughter cells predominantly bud in the same direction of their mother (nematic order parameter, S = 0.900), at approximately 180 degrees, such that they form loosely linear chains of spherical cells. This high level of directionality is reflected in a high nematic order parameter. This analysis quantifies 148 budding events from 28 colonies chosen at random. B’) Simulated angular distribution of *K. petricola*. If growth is random (isotropic), we expect an even distribution of budding angle, which is reflected in the lower nematic order parameter of 0.313. Our model simulated the growth of 50 colonies starting from a single cell for 36 hours. C. DNA staining in fixed cells. Cells are uninucleate. Scale bar, 10 μm. D. Time until a newly budded cell begins producing its first daughter. Cells are capable of a remarkably quick turnaround time of 10 min, but could also take as long as 320 min (N = 28, mean = 110, SD = 64.6, CV = 58.7%). E. Cell cycle length (min). This is a measurement of the full cell cycle duration. Cell cycle length is highly variable (N = 28, mean = 499, SD = 121, CV = 24.3%).

We evaluated if indeed these division and colony patterns were non-random by simulating colony patterns based on random division directions. While the orientation of new buds in the actual cells is highly uniform and often opposite from its mother’s birth site (Figure 4B), the orientation of simulated random budding shows that a highly variable distribution of orientations is possible (Figure 4B’). Of 152 budded cells observed, only 8.6% budded in a direction other that opposite the birthsite. Buds appear to grow uniformly and each bud receives a nucleus (Figure 4C). Cell cycle lengths are highly variable with a mother initiating a new bud in as little time 10 min, although the average time to start a bud was 110 min (SD = 64.6 min, CV = 58.7%, Figure 4E-F). Thus, this system is remarkable for using highly spherical cells to produce a polarized colony shape akin to filamentous growth but generated through buds rather than hyphae.

### Simultaneous production of multiple buds per cell cycle in *Aureobasidium pullulans*

*A. pullulans* was isolated from several locations including both plankton tow sites, sponge from the Great Harbor and sediment from Sippewissett Marsh. It is a black yeast species known for its potential biotechnological significance as a producer of pullulan (poly-alpha-1,6-maltotriose), a biodegradable, extracellular polysaccharide [64], and it encodes enzymes capable of plastic degradation [48, 65, 66]. This species is globally ubiquitous and has been isolated from osmotically stressed environments, such as salterns, rocks, and glacial and subglacial ice [67]. A major feature of budding yeast morphogenesis, observed not only in *S. cerevisiae* but also *C. albicans* and the distantly related basidiomycetes *C. neoformans* and *U. maydis*, is that only a single bud is produced each cell cycle. Singularity in budding ensures that with each mitosis, a daughter and mother each receive a single nucleus. Remarkably, however, the yeast form of *A. pullulans* produced multiple buds simultaneously, up to 6 from the same vicinity of a mother cell. The number of buds a mother produced could vary each cell cycle, but a single mother displayed a tendency to produce similar numbers of buds in consecutive division cycles, and the sites of bud emergence were often re-used (Figure 5A, Videos S3 and S4). Almost 80% of scored cells produced >1 bud, and of those, 64% produced multiple rounds of simultaneous buds (Figure 5B). Mean cycle length was 159.5 min and was quite variable (N = 82, SD = 80.5 min, CV = 50.5, Figure 5C), but cell cycle duration was not obviously correlated with the number of buds a mother produced in a given cell cycle (Figure 5C’).

**Figure 5.**
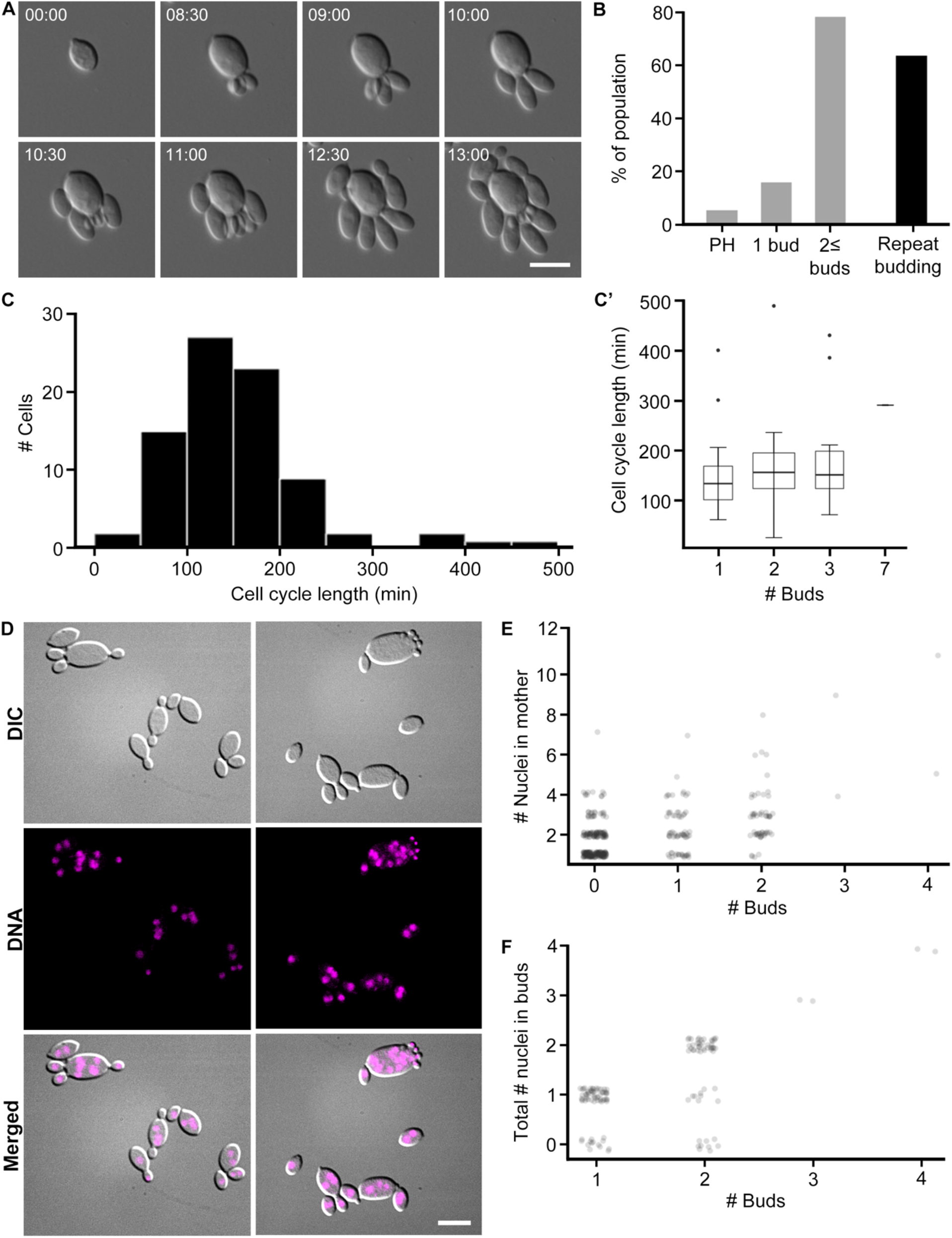
*A. pullulans* is multinucleate and is capable of producing multiple buds simultaneously. A. Representative DIC images of *A. pullulans* cell growth and development at 40x DIC. Cells may produce multiple buds at once, and sometimes do so in quick succession and from the same site. Scale bar, 10 μm. Time, Hours:Minutes. B. Percentage of population that produce multiple buds. In grey are the percentage of cells that are become pseudohyphal and produce one or multiple buds (N =75). 59 cells (79%) produce multiple simultaneous buds. Of those, 38 cells (64%) repeatedly produce multiple simultaneous buds, indicated by the black bar. C. Cell cycle length (min). This is a measurement of the full cell cycle duration (N = 82, mean = 159.5 min, median = 150, SD = 80.5, CV= 50.5). C’) Cell cycle length based on number of buds that a mother produces. The number of simultaneous daughter cells that a mother produces does not seem to affect the length of its cell cycle. Box widths are representative of sample size. D. DNA staining in fixed cells reveals that mother cells are multinucleate. Buds are predominantly uninucleate, but some appear to have no nuclei. Scale bar, 10 μm. E. Number of nuclei in mother cells binned by the number of buds it produces. This counts only the nuclei that are in the mother cells (N = 371). Data was scored from fixed cells. Mothers that produce more buds appear to have more nuclei. F. Total number of nuclei in buds. This counts the number of nuclei in buds from the same mother (N = 127 family groups). Data was scored from fixed cells. While this does not explicitly reflect the number of nuclei in each bud, we observed no multinucleate buds.

We speculated that the variable numbers of buds might reflect a variable number of nuclei in the mother cell. Mother cells did have highly variable numbers of nuclei based on nuclear staining of fixed cells (Figure 5D). Many multinucleate mothers, particularly those that gave birth to repeated generations of multiple buds, were quite large and irregularly shaped compared to their progeny (Video S3). Mothers with the most nuclei tended to produce higher numbers of buds in a given division cycle, but the correlation was weak (Figure 5E). Buds generally received no more than one nucleus each, and in some cases the nucleus was deposited even when buds are extremely small (Figure 5D-F). Thus, mothers appear to produce some number of nuclei in the absence of budding, and then segregate nuclei to nascent buds. How nuclear division and budding are coordinated will require analysis of nuclear dynamics in live cells. Thus, this system presents multiple surprises including the ability of cells to host several coexisting polarity sites, and to populate daughter cells with a single nucleus despite a reservoir of many nuclei from which to choose.

### Meristematic divisions, filamentation and cellularization in *Phaeotheca salicorniae*

*P. salicorniae* was isolated twice from plankton tows, once in each of our main tow areas. The genus Phaeotheca was first identified in 1981 [68]. *P. salicorniae* was first identified in 2016 and thus has a very limited literature [69]. We also isolated the very closely related *Phaeotheca triangularis* [70] from both coral and sponge. We focused on *P. salicorniae*, which has the most complexity in morphology and division patterns and is least like any of the well characterized model yeasts or filamentous fungi in our study. In *P. salicorniae* we have identified yeast cells and hyphal cells, with cells able to switch bidirectionally between these two types. Colony growth begins from a yeast cell (Figure 6A’, Video S5) which swells and elongates slightly and then goes through a series of divisions by forming septa. The first septum is placed in the center of a slightly ovoid cell. Subsequent divisions are perpendicular to the previous division plane, producing a small cluster of cells with shapes reminiscent of sections of a pie (Figure 6A’’). After this, cells transition from this small group of 6-8 or more cells separated by septa to a small cluster of cells that appear to emerge as spherical cells within the cluster. This is termed meristematic conversion and these cells appear to be embedded in some sort of extracellular matrix that can be rich in melanin.

**Figure 6.**
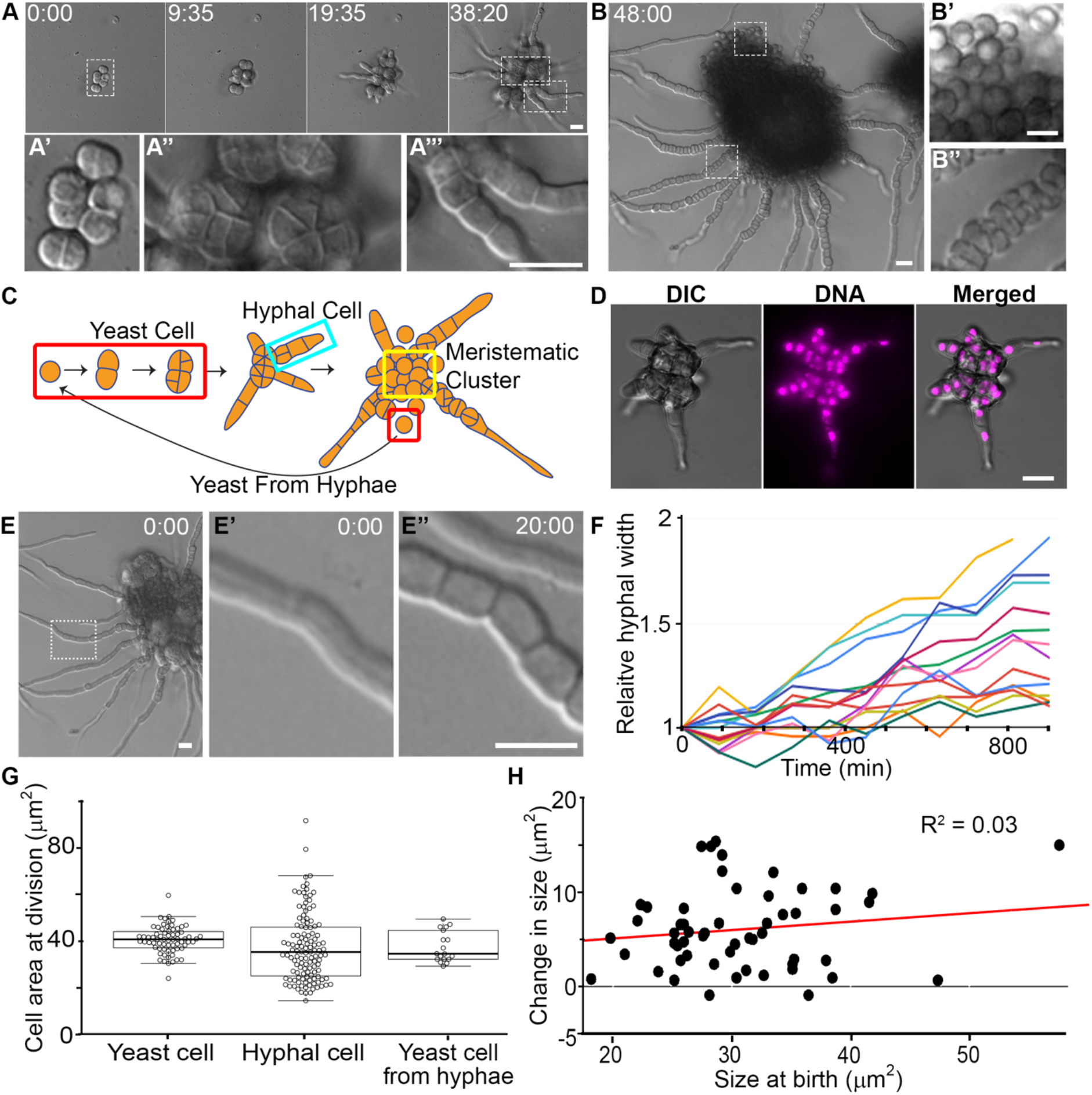
*P. salicorniae* colony growth involves meristematic division and filamentation. A. Representative DIC micrographs of *P. salicorniae* colony growth over the course of 38 hr. Insets show different features of the colony A’ yeast cell, A’’ wedge shaped cells, A’’’ hyphal cells. Scale bar, 10 μm. Time, Hours:Minutes. B. Representative DIC micrograph of a *P. salicorniae* colony 48 hr after the start of growth. Insets (B’ B’’) show different cell types within the colony. Scale bar, 10 μm. Time, Hours:Minutes. C. Model for the growth of a *P. salicorniae* from a single colony. Initial growth begins with a yeast cell. Growth is isotropic and divisions bisect the cell resulting in wedge shaped cells (see A’’). At a critical size some yeasts cells begin to grow hyphae. Hyphal cells continue to divide until eventually the hyphae bursts with cells that are morphologically similar to the initial yeast cell. D. DNA staining in fixed cells. *P. salicorniae* compartments contain only a single nuclei. Representative image of a *P. salicorniae* colony in DIC, stained with DAPI, and in merge. Scale bar, 10 μm. E. *P. salicorniae* hyphae continue grow wider while elongating. A representative image of a colony at the start of filming. Inset (E’, E’’) showing hyphal width at start of filming (0:00) and the conclusion of filming (12:00). Scale bar, 10 μm. Time, Hours:Minutes F. Quantification of the change in relative hyphal thickness from the start to end of filming. Each line represents an individual hypha, all hyphae increase in thickness over the length of filming. G. Quantification of cell size at division in various cell types. The mean size at division is not significantly different between cell types (p > 0.05, t-test) but hyphal divisions are significantly more variable than yeast cells exhibiting both much larger and much smaller cells (p < 0.01, f-test). H. Quantification of the cell size at birth vs. the cell size at division for hyphal cells and yeast cells.

Two events occur in this meristematic colony. Some cells transition into a second cell type by producing hyphae which extend by polarized growth (Figure 6A’’’). At the same time, an abundance of yeast-like cells that are similar in appearance to the original cell (Figure 6B’) are produced and remain associated in a matrix (Video S6). Eventually these yeast cells are released by the breakdown of the matrix around them. In the hyphae, cells proximal to the colony septate, swell, and produce new spherical cells within the hyphae (Figure 6B’’, Video S6). These spherical cells appear to be similar to the original yeast cells and can form multiple, orthogonal septa while inside the hyphae. This suggests the ability to switch from hyphal back to yeast-like growth from within a hypha, although it could represent the formation of spores rather than vegetative yeast. As the hyphae grow, the remaining yeast cells continue to divide meristematically and the septa are less clearly visualized (Figure 6C).

In addition to alternating between polar, hyphal growth and isotropic, yeast-like growth, we found several other characteristics of this fungus that differ from conventional growth modes. First, the hyphae of *P. salicorniae* contained only one nucleus per compartment (Figure 6D). Second, *P. salicorniae* hyphae did not have a fixed width and continued to expand laterally while elongating (Figure 6E-F). We also found that yeast and hyphal cells appeared to have different degrees of cell size control. Both the yeast and the cells within the hyphae divided with a similar mean area of 40.4 μm^2^ and 37.5 μm^2^, respectively (not significant, P > 0.05, t-test, N > 62) (Figure 6G). However, the size at division was much more variable in the hyphal cells, which had a standard deviation of 15.9 μm^2^, compared to yeast cells with a standard deviation of just 5.8 μm^2^ (p < 0.01, F-test, N > 62). Consistently, in hyphal cells, the change in size is virtually uncorrelated with the size at birth, indicating a lack of size control (Figure 6H). Thus, this system shows striking orthogonal divisions unlike those of any model fungal system studied to date for cytokinesis, hyphal growth at the colony periphery, and striking alternation in cell types.

## Discussion

Very little is known regarding the distribution, diversity and morphogenesis of marine-derived fungi. Furthermore, there appears to be strong potential for some fungal species to be amphibious as displayed by their highly adaptive nature and their isolation from both terrestrial and marine environments [71–73]. The goal of this study was to identify how fungi found in the marine environment may grow and divide under different environmental challenges faced in the ocean.

Despite being geographically, temporally and methodologically limited, this study captured substantial fungal diversity. This is surely a significant undersampling of diversity as we used only limited culture conditions, and species which require very specific nutrient requirements were likely missed in our pipeline. A culture-based approach was taken because the goal of this study was to analyze morphogenesis and division of cells. This has the advantage that these systems can now be carefully studied in the lab and empirically analyzed. We are still in the process of identifying other isolates from our collection, but several species including *H. werneckii* appeared several times and in several places, suggesting that at least some of the species are reasonably abundant. Interestingly, 25% of our 77 sequenced isolates were Basidiomycetes whereas only 10, or 2.15%, of the 465 marine isolates identified by Shearer and colleagues [71] were of that phylum. This is certainly not an exhaustive analysis of the fungal diversity present in this location. Metagenomic analysis of samples in the future will give a more comprehensive snapshot at the actual fungal diversity.

There were substantial differences in growth and division amongst the four black yeasts analyzed both between one another and in comparison to well characterized yeast model systems. These differences may support these black yeasts’ flexibility and ability to exist under the challenges associated with marine environments. Remarkably, the two distinct modes of division in the model yeasts-fission and budding-coexist in the isolates of *H. werneckii*. In this system, cells systematically alternate between division by fission and budding, readily transitioning between them in a single generation. It is unclear what, if any, adaptive advantage this toggling between forms provides these cells. One possibility is that it represents a form of bet hedging. Depending on specific environmental stress conditions, budding or fission may be more advantageous, and by alternating between these states on a short timeframe, some subset of the population will be ready for any rapid challenge.

A second surprise came from *A. pullulans* which simultaneously produced up to six buds from a multinucleate mother cell. We also isolated the closely related black yeast, *Aureobasidium melanogenum*, once from coral and once from sponge from Great Harbor. *A. melanogenum* also exhibited the pattern of producing multiple buds from the same site, although we did not carry out a detailed analysis. This multi-budding phenotype is in striking contrast to the tightly controlled single bud per cell cycle produced by *S. cerevisiae*. Singularity in budding is a mechanism to coordinate nuclear division and morphogenesis to ensure that each daughter cell receives only one nucleus. In *A. pullans* it is unclear what determines which cells in the population ultimately become mother cells, as the buds produced are generally much smaller and more homogeneous in size that the mothers, which are often highly irregular in shape and size. How the nuclear cycle of the mother cells is coordinated with the multi-budding program, how specific nuclei are selected for deposition into buds, and how it is ensured that each bud receives a nucleus remain fascinating problems for future study in this system.

Yet another striking growth pattern that is distinct from the model yeasts was seen in *K. petricola* which produced spherical cells that appeared to expand isotropically. These bubble-like cells generated linear chains at the colony level. The chains displayed a highly non-random budding pattern although on occasion, the chain will sprout a chain through a mother producing a second bud at a distinct orientation. This allows these spherical cells to generate dendritic colonies. We speculate that despite the very isotropic growth, there is some mark at the site of bud initiation that persists and biases the site of the next bud. Testing this hypothesis must await the development of molecular tools for this system.

Finally, the most complex isolate was *P. salicorniae*, a very recently discovered species with limited descriptions in the literature. The myriad of growth patterns seen in this organism suggest levels of developmental control more complex than in the other black yeasts. The different cell types and meristematic growth resemble other species in the genus such as *Phaeotheca triangularis*, but what stands out as remarkable compared to many traditional fungal models are the series of orthogonal divisions that lead to triangular-shaped compartments and the intercalary growth of hypha. How division planes are positioned and which cells are fated for hyphal growth from the meristematic colony are fascinating open questions from this new system.

These studies reveal the morphological plasticity present within individual fungal species and the limited view that study of a few model systems have presented as to the rules governing fungal shape and division. Unlike many of the model systems cultivated in lab, there is little evidence of tight size control over the division cycle for several of the marine-derived yeasts, with high degrees of variability in cell size for 3 of the species we examined. This highlights the potential of observing diverse non-model biological systems to test our deeply held models of biological phenomena. We predict many fundamental biological mechanisms have higher degrees of plasticity than appreciated from studies limited to model systems.

## Methods

### Collection

#### Environmental ocean samples

We collected 17 environmental ocean water samples on eight days spanning from summer 2016 to winter 2018 (Figure 1A). Only one sample was collected each during summer 2016 and winter 2018; the majority were collected during summer and fall 2017. We used a plankton net that was 1 m in diameter with a 504 μm mesh. This was trailed for 10 min behind a Boston whaler at an average over-ground speed of 1.2 m/s. The top of the net was kept just under the water surface. Tows were 1-1.5 km offshore and average flowthrough volume was 2400 m^3^.

#### Sediment samples

Coastal sediment samples were collected nearshore using sterile 50 ml conical tubes (this always included some amount of water as well). Samples were collected from beaches adjacent to Buzzards Bay and Martha’s Vineyard Sound. These sites included Trunk River, Surf Drive Beach, Fay Beach, Garbage Beach, Dog Beach, Quissett Harbor, the Knob, Sippewissett Marsh, and by the Woods Hole Town Dock in Great Harbor (Figure 1A).

#### Coral and sponge samples

Coral and sponge samples were collected off Garbage Beach in Woods Hole, MA during July and August 2018 and February 2019. Samples were collected 9-13 m below sea level. Corals were rinsed with sea water filtered through a 0.22 μm filter and removed from their substrate of rock or sponge. For each coral sample, an average of 42 polyps of the same color from the same colony were ground up using a mortar and pestle with filtered seawater added with a ratio of 1 ml water to 1 g of sample. Additionally, some coral and sponge were inhabited by small black worms, which we removed as much as possible prior to grinding. Average mass per sample was 7.74 g. 100 μl of liquid supernatant and 150 μl of slurry (solid ground-up coral) were plated in duplicate on YPD, MEA, and PDA saltwater plates. Sponge prepared in the same manner and water from the same location was a control.

Polyps without Symbiodinium (aposymbiotic, from February collection) are paler in color and polyps with Symbiodinium (symbiotic) are darker and browner [74]. We observed a range of colors including white and light pink to darker purple and brown. Some colonies contained polyps of multiple different colors.

### Media

Samples were plated on three types of media to offer different carbon sources; Yeast Extract (YPD; yeast extract [BD #212720] 10 g/L, peptone [BD #211677] 10 g/L, dextrose [VWR chemicals #BDH9230] 20 g/L, agar [BD #214010] 12 g/L), Malt extract (MEA; malt extract [BD #218630] 20 g/L, peptone 6 g/L, dextrose 20 g/L, agar 12 g/L), and potato dextrose (PDA; potato dextrose broth [Sigma Aldrich #P6685] 24 g/L, agar 12 g/ L). All media were made with filter sterile seawater, or 36 g/L of Instant Ocean (Spectrum Brands, Blacksburg, VA) if plates were made inland without easy ocean access. Antibiotics were added to select for fungal isolates (carbenicillin 100 μl/ml, chloramphenicol 25 μl/ml and tetracycline 10 μl/ml).

### Filming

We collected long-term (up to 36 hr) DIC time-lapse data with a Nikon Eclipse Ti2 inverted microscope with a perfect focus system using a 40x dry objective lens using a Nikon DS-Qi2 camera. Samples were imaged on agar pads made of the same media type on which they were originally isolated. We imaged up to six isolates (one agar pad per isolate) with four areas of interest each per filming session.

Agar pads were made by melting agar media and pipetting the liquid into a mold. The mold was constructed by sandwiching a 1/32 in rubber sheet cut like a rectangular frame missing the top side (so the mold may be filled in) sandwiched between two glass microscope slides and held together with binder clips. The liquid was let to cool and solidify for 15 min. Pads were cut with sterile razor blades to be approximately 1 cm^2^. In the following order, we placed on a 28×65 mm coverslip: 6 μl of dilute liquid culture, the agar pad, 2 μl liquid media and lastly the microscope slide. The coverslip, agar pad and microscope slide sandwich is then sealed with VALAP. Several holes were created in the VALAP between the coverslip and slide with a 27 gauge needle to allow oxygen flow. VALAP is **Va**seline, **La**nolin [Sigma-Aldrich #L7387], and **P**araffin [Sigma-Aldrich #327204] in a 1:1:1 mass ratio.

### DNA Staining

H. werneckii was grown in liquid culture for 18 hr at 30 °C in liquid media (YPD + 0.6 M NaCl). Cells were fixed with 3.7% formaldehyde and 25 mM Dithiothreitol (DTT) (US Biological #D8070) for 1 hour. Cells were washed with 1x phosphate buffered saline (PBS) 3 times, then stained with Hoechst 33342 (Invitrogen #H3570) in PBS for 30 min.

K. petricola was grown in liquid culture for 92 hr at 22 °C. Cells were fixed in 100% methanol on ice for 20 min, washed with 1x PBS, then stained with NucBlue Live ReadyProbes (Thermofisher #R37605) in PBS.

A. pullulans was grown in YPD liquid culture for 24 hr at 24 °C with Complete Supplement Mixture (CSM) (Thermofisher #A13292) and 2% Dextrose + 0.5M NaCl. Cells were fixed in 70% ethanol on ice for 10 min, washed with 1x PBS + 0.5 M NaCl, then stained in 1x PBS + .5M NaCl with 50 ng/mL 4’,6-diamidino-2-phenylindole (DAPI) (Sigma-Aldrich #D9542).

P. salicorniae was grown affixed to slides coated with Concanavalin A (Sigma-Aldrich #L7647) immersed in liquid media (YPD + 0.6M NaCl) for 48 hr at 22 °C. Cells were fixed on the slide with 100% ice cold methanol for 20 min, washed with 1x PBS, then stained with Hoechst 33342 in PBS.

### Quantifying cell characteristics

Cell size metrics were measured using ImageJ [75]. Area was measured by tracing the outline of a cell with the freehand selection tool. Length and width were measured using the line measurement tool. To measure the time between divisions, the number of frames between two events was counted and then multiplied by the frame rate.

### Modeling colony growth of *K. petricola*

Single *K. petricola* cells were imaged for 36 hr to observe their growth into a colony using a DIC microscope with 40x magnification acquiring images every 5 min. The resulting movies were segmented using the Empirical Gradient Threshold method for DIC images [76]. The segmentation could not identify individual cells but performed well in distinguishing the cell colony from the background and resulted in a binary image, or mask, where pixels were set to 1 if they were part of a cell (colony) and 0 otherwise. In the following data analysis, we aimed to parametrize growth of the cell colony and the overall directionality of the colony growth.

Since we only considered 2-D cell colonies, i.e. ones that grew flat on the cover slip, we used the area covered by cells as a measure for growth, which is determined by the number of masked pixels. In order to assess the directionality of growth, we first computed the differential segmented image between each frame and the previous one, resulting in a binary image that measures growth between acquired frames. We then calculated the center of mass (COM) of the segmented first frame of each movie and established an arbitrary reference line across the COM. For each pixel in the differential segmented images, we calculated the angle between this reference line and the line connecting the pixel and the COM. The distribution of these angles holds information about the growth direction in each frame. In order to parametrize this angular distribution, we adjusted the reference line such that the mean angle is 0° and calculated the nematic order parameter averaged over all pixels: 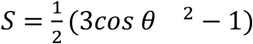. As a result, we can follow the directionality of the colony growth throughout the entire movie, through the nematic order parameter which in our case takes values between 1 (anisotropy) and 0 (isotropy).

A meaningful interpretation of the nematic order parameter in our context was not possible without any reference values. We therefore formulated the null-hypothesis that growth of a cell colony is isotropic and no direction is preferred. Based on this assumption, we simulated segmented movies that were analyzed in the same way as the experimental data and yielded the expected nematic order parameter for isotropic growth close to 0. The simulation was rather simple, i.e. individual cells did not grow but were introduced from one frame to the other with their full size. Budding times and cells size were drawn randomly from a pool of values determined manually from the experimental data beforehand.

### Molecular identification of isolates

Cells were grown on the same type of media from which they originated. Yeast were harvested from agar plates while filamentous fungi were harvested from liquid media. We utilized the DNeasy Plant Mini Kit (QIAGEN #69104) to extract DNA as per the manufacturer’s protocol. For filamentous isolates for which the DNeasy Plant Mini Kit did not work well, we performed a phenol chloroform extraction (modified from [77]). All samples were amplified using the ITS1, ITS2, D1/D2, and V9 rDNA regions for molecular identification (Table S1). The RPB2 gene was additionally sequenced when rDNA failed to distinguish between closely related species, such as *A. pullulans* and *A. melanogenum*.

### Isolation of Genomic DNA using Phenol/chloroform

Filamentous fungi were grown in liquid YPD for approximately a week on a rotator at 20 °C. Colonies were removed with a sterile spatula and placed on a stack of sterile paper towels to wick off liquid. Dried colonies were transferred to an Eppendorf tube, with 750 μl of lysis buffer (50 mM Tris-HCl of pH 7.4, 50 mM EDTA, 3% SDS and 1% 2-mercaptoethanol) and vortexed. Then incubate at 65 °C for 1 hr. 750 μl of phenol/chloroform/isoamyl alcohol (Sigma-Aldrich #77617) was added to the tube and then briefly vortexed. The tube was spun at 12,000 x g for 10 min and the aqueous layer was removed to a new tube. A second phenol/chloroform extraction was then performed on the aqueous phase. Following the second extraction, the aqueous phase was transferred to new tube and 20 μl of 3 M NaOAc and 700 μl isopropanol precooled to −20 °C was added. The tube was inverted gently to mix then spun at 12,000 x g for 30 sec to pellet DNA. The supernatant was removed and discarded. The pellet was resuspended in 300 μl of TE (100 mM Tris-HCl, 10 mM EDTA, pH 8.0) at 65 °C for 15 min followed by finger vortexing. DNA was then precipitated by the addition of 10 μl of 3 M NaOAc and 700 μl ethanol at −20°C. The tube was spun at 12,000 x g for 2 min and the supernatant was removed. The pellet was washed with 500 μl of 70% ethanol at −20°C. The pellet was then spun at 12,000 x g for 30 sec the supernatant was removed and the pellet was allowed to air dry at room temperature for 30 min. The pellet was then resuspended in 100 μl of sterile DI water.

## Acknowledgements

CMF, ASG, and LMYM-F thank the MBL for support including Scott Bennett and the MBL Marine Resources Center for help in collecting specimens, Loretta Roberson at MBL for advice on working with coral, and the Physiology and Microbial Diversity courses at MBL for collaborative interactions. CE was a student Physiology course participant. We also thank Nikon at MBL and the Nikon Imaging Center at HMS for microscopes and imaging advice. LMYM-F thanks the Springer Lab at HMS for initial help with DNA extraction.

## Author Contributions

LMYM-F, CMF and ASG conceived of and designed the collection. LMYM-F performed the majority of collecting, culturing of samples, and imaging of isolates. ASG did the majority of writing with contributions from LMYM-F, JMV-M, BMS, EJDV and CMF. LMYM-F and JMV-M performed the DNA extraction and species identification. LMYM-F created Figure 1. BMS created Figure 2. JMV-M and LMYM-F performed experiments and analysis in Figure 3. CE conceived of the method for analysis of colony morphology in Figure 4. LMYM-F performed the colony morphology analysis and did the quantitation in Fig 4D and E. EJDV performed the DNA staining in Fig 4D. CMF and LMYM-F performed the imaging and analysis in Figure 5A-C. SVD and DJL performed the DNA imaging and analysis in Figure 5D-F. BMS performed the DNA imaging and analysis in Figure 6. Table 1 created by JMV-M.

## Supplemental Materials

Video S1. DIC timelapse of *H. werneckii* (40x). Scale bar, 10 μm. Time, Hours:Minutes.

Video S2. DIC timelapse of *K. petricola* (40x). Two examples of colony growth. Scale bar, 10 μm. Time, Hours:Minutes.

Video S3. DIC timelapse of *A. pullulans* (40x). This mother cell repeatedly produces multiple simultaneous buds from the same site. Scale bar, 10 μm. Time, Hours:Minutes.

Video S4. DIC timelapse of *A. pullulans* (40x). 3 more examples of budding mother cells demonstrating the variability of cell morphology. Scale bar, 10 μm. Time, Hours:Minutes.

Video S5. DIC timelapse of *P. salicorniae* (40x). Growth of initial yeast cells and sprouting of hyphae which then develop septa. Scale bar, 10 μm. Time, Hours:Minutes.

Video S6. DIC timelapse of *P. salicorniae* (20x). At left, overview of later stages of colony growth. At right, close-ups of hyphae septating and reverting back to yeast cells. Scale bars, 10 μm. Time, Hours:Minutes.

Table S1. Primers used for DNA sequencing.

