## Supplementary material for "Unconventional cell division cycles from marine-derived yeasts": Table S1

Table S1. Primers used for DNA sequencing.

| **Primer** | **Sequence** | **Region** |
|---|---|---|
| ITS1 | TCCGTAGGTGAACCTGCGG | |
| ITS4 | TCCTCCGCTTATTGATATGC | rDNA ITS1 |
| 5.8S-F | GTGAATCATCGARTCTTTGAAC | |
| 28S1-R | TATGCTTAAGTTCAGCGGGTA | rDNA ITS2 |
| NL-1 | GCATATCAATAAGCGGAGGAAAAG | |
| NL-4 | GGTCCGTGTTTCAAGACGG | rDNA D1/D2 |
| 1389F | TTGTACACACCGCCC | |
| 1510R | CCTTCYGCAGGTTCACCTAC | rDNA V9 |
| RPB-PenR1 | GTTCACDCAACTYGTGCGYGA | |
| RPB-PenR2 | GGCAGGGTGAATYTCGCAATG | RNA Polymerase II (RPB2) |
